# Suppressor of Fused controls perinatal expansion and quiescence of future dentate adult neural stem cells

**DOI:** 10.1101/454751

**Authors:** Hirofumi Noguchi, Jesse Garcia Castillo, Kinichi Nakashima, Samuel J. Pleasure

**Author notes:** Corresponding authors: Samuel J. Pleasure, MD/PhD, Department of Neurology, University of California, San Francisco, 675 Nelson Rising Lane Mail Stop #3206, San Francisco, California 94158, University of California, San Francisco.

## Abstract

Adult hippocampal neurogenesis requires the quiescent neural stem cell (NSC) pool to persist lifelong. The establishment and maintenance of quiescent NSC pools during development is not understood. Here we show that Suppressor of Fused (Sufu) controls establishment of the quiescent NSC pool during mouse dentate gyrus (DG) development through regulation of Sonic Hedgehog (Shh) signaling. Deletion of *Sufu*in NSCs at the beginning of DG development decreases Shh signaling leading to reduced proliferation of NSCs, resulting in a small quiescent NSC pool in adult mice. We found that putative adult NSCs proliferate and increase their numbers in the first postnatal week and subsequently become quiescent towards the end of the first postnatal week. In the absence of Sufu, postnatal expansion of NSCs is compromised, and NSCs prematurely become quiescent. Thus, Sufu is required for Shh signaling activity ensuring the expansion and proper transition of NSC pools to quiescence during DG development.

## Introduction

Newborn neurons are generated in two restricted regions of the adult rodent brain: the cortical subventricular zone (SVZ) and the dentate subgranular zone (SGZ) (Altman and Das, 1965; Eriksson et al., 1998; Kuhn et al., 1996; Lois and Alvarez-Buylla, 1993). Adult neurogenesis in the dentate gyrus (DG) has been implicated in hippocampal-dependent memory and learning (Deng et al., 2010). Newly generated neurons produced from neural stem cells (NSCs) residing in the SGZ are constantly added to the granule cell layer (GCL) and integrated into the existing hippocampal circuitry (Imayoshi et al., 2008). Persistence of adult hippocampal neurogenesis relies on the proper maintenance of NSCs even after development. However, little is known about the developmental programs governing the production and maintenance of long-lived NSCs.

Quiescence of NSCs during early development has been proposed to play a key mechanism for maintaining the NSC pool throughout life (Furutachi et al., 2013; Kawaguchi et al., 2013; Mira et al., 2010; Song et al., 2012). NSCs enter a quiescent state in a spatiotemporal manner during development, and this step is critical for ensuring the appropriate sized NSC pool for adult neurogenesis. At the beginning of forebrain development, NSCs are highly proliferative, but gradually lose proliferation competence with development and enter a quiescent state (Furutachi et al., 2015). The failure to transition to a quiescent state during development triggers continuous proliferation of NSCs and leads to premature exhaustion of the NSC pool (Furutachi et al., 2015). Furthermore, the NSC pool must be properly established since NSCs can only undergo a limited number of rounds of cell division prior to terminal differentiation. Live imaging of NSCs in adult DG and thymidine-analog based cell tracing analysis demonstrated that quiescent NSCs undergo a series of asymmetric divisions to produce neurons, and subsequently are consumed by symmetric differentiation into astrocytes or neurons (Calzolari et al., 2015; Encinas et al., 2011; Pilz et al., 2018). Indeed, adult neurogenesis and NSC pool have been shown to decline with aging (Kuhn et al., 1996; Lugert et al., 2010), suggesting that for neurogenesis to persist throughout life, the size of putative quiescent NSC pool must be established during development before the NSCs transition to a quiescent state.

Long-lived NSCs in the DG originate from the ventral hippocampus at E17.5 (Li et al., 2013). These cells migrate along the longitudinal axis of the hippocampus from the temporal to septal poles and eventually settle in the ventral and dorsal DG. The initial production and maintenance of long-lived NSCs is dependent on Shh signaling and Shh ligands, produced by local neurons in the embryonic amygdala and the postnatal DG (Li et al., 2013). Blocking Shh signaling by deleting *Smoothened (Smo)* from responsive cells in the DG or ablation of Shh ligands from local neurons impairs the emergence of long-lived NSCs and results in diminishing the NSC pool (Han et al., 2008; Li et al., 2013). These findings highlight the significance of Shh signaling in production of the NSC pool during development. What is not clear yet from these studies is how Shh signaling activity is spatiotemporally regulated to ensure the expansion of the NSC pool during DG development and the role of Shh signaling in the transition of NSCs to a quiescent state.

Shh signaling is critical at early stages of embryonic brain development. Thus, complete ablation of Shh signaling activity by *Smo* deletion or the constitutive activation of Shh signaling by expressing an active Smo mutant (SmoM2) severely compromise the initial steps of DG development (Han et al., 2008). The embryonic nature of this phenotype prevents the further analysis of specific roles of Shh signaling in postnatal DG development, particularly in the production and maintenance of postnatal NSCs. To circumvent this, we are utilizing a Cre-loxP based system that allows spatiotemporal analysis of Shh signaling activity by genetic manipulation of the Shh signaling inhibitor, Suppressor of Fused (Sufu), a Gli-binding protein with an indispensable role in embryonic development. Conditional deletion of *Sufu* in a spatiotemporal manner allowed us to examine how uncontrolled Shh signaling influences various aspects of NSC behavior during DG development. Our earlier studies showed that Sufu is important for the specification of NSC fate decision during cortical development via regulating Shh signaling activity (Yabut et al., 2015). In this report, we set out to determine the contribution of Sufu in regulating Shh signaling during DG development and how Sufu and Shh signaling are involved in the mechanisms governing the expansion of long-lived NSCs and their transition to the quiescent state during DG development. Intriguingly, we find that deletion of *Sufu* decreases Shh signaling in NSCs during DG development – this is in distinction to the situation in the neocortex where loss of *Sufu* increases Shh signaling. Long-lived NSCs expand in the early part of first postnatal week, but proliferation of these NSCs is impaired in the absence of Sufu, resulting in a decreased NSC pool in the adult DG. We also found that long-lived NSCs gradually become quiescent towards the end of first postnatal week. However, *Sufu* deletion precociously triggers this transition to the quiescent state. Taken together, these results indicate that Sufu, through regulation of Shh signaling activity, is important for ensuring the expansion of long-lived NSCs and the timely transition to a quiescent state during DG development.

### Materials and methods

#### Animals

Mice carrying the floxed Sufu allele (Sufu^fl^) were kindly provided by Dr. Chi-Chung Hui (University of Toronto) and were genotyped as described elsewhere (Pospisilik et al., 2010). The following mouse lines were obtained from Jackson Laboratory (Bar Harbor, Maine): Gli1^CreERT2/+^ (stock 007913), Gli1^LacZ/+^ (Stock #008211), Rosa-AI14 (Stock #007908), SmoM2 (Stock #005130), hGFAP-Cre (Stock #004600). Both male and female mice were analyzed with no distinction. All mice used in this study were maintained on a 12-h light/dark cycle with free access to food and water. The day of vaginal plug was considered embryonic day 0.5. Mouse colonies were maintained at University of California San Francisco (UCSF) in accordance with National Institutes of Health and UCSF guidelines.

#### Tamoxifen and Thymidine analog administration

Tamoxifen (Sigma) was dissolved in corn oil at 10 mg/ml. Pregnant mice were intraperitoneally administered 2 mg of tamoxifen with 27-gauge needles. For bromodeoxyuridine (BrdU) labeling, mice were subcutaneously (neonatal pups) or intraperitoneally (adult mice) injected with BrdU (Sigma) dissolved in saline (0.9% NaCl) at a dose of 50 mg/kg. For two thymidine analog labeling, chlorodeoxyuridine (CldU, Sigma) dissolved in saline was subcutaneously injected into neonatal pups at a dose of 42.5 mg/kg followed by a single intraperitoneal injection of 57.5 mg/kg iododeoxyuridine (IdU, Sigma) in 0.2 N NaOH 0.9% NaCl solution for five days at 8 weeks old.

#### Tissue preparation

To prepare embryonic brain tissue, pregnant mice were sacrificed on the indicated developmental day, and embryos were perfused successively with phosphate-buffered saline (PBS) and ice-cold 4% paraformaldehyde (PFA) in PBS, pH 7.2. For preparation of postnatal and adult brains, pups and adult mice were deeply anesthetized before perfusion with 4% PFA in PBS. Brains were dissected and postfixed with 4% PFA in PBS overnight at 4°C. For cryoprotection, fixed brains were stored in 30% sucrose in PBS at 4°C. The brain was embedded in optimal cutting temperature (OCT) compound (Tissue Tek, Sakura Finetek, 25608-930) and frozen at −80°C for cryosectioning. Frozen brains were serially sectioned with Leica CM 1850 (Leica Microsystems, Wetzlar, Germany) in the coronal or sagittal plane at 16 μm thickness. Every fifteenth sections were serially mounted on individual Colorfrost Plus Microscope Slides (Fisher Scientific) in order from anterior to posterior (coronal section) or medial to lateral (sagittal section), and preserved at −20°C until use.

#### LacZ Staining and In situ hybridization

Animals for LacZ staining were perfused with 4% paraformaldehyde (PFA) and the dissected brains were postfixed with 4% PFA for 2 hr at 4°C. Cryosections were washed with PBS, and X-gal staining was developed at 37°C overnight in the staining solution (5 mM K_3_Fe(CN)_6_, 5 mM K_4_Fe(CN)_6_, 5 mM EGTA, 0.01% deoxycholate, 0.02% NP40, 2 mM MgC1_2_, and 1 mg/ml X-gal). Sections were postfixed with 10% formalin at room temperature overnight, followed by counterstain with nuclear-fast red (H-3403, Vector Laboratories) at room temperature for 10 min before proceeding for dehydration (70%, 95%, 100% ethanol, xylene twice) and coverslipping with Mount-Quick (Ted Pella).

#### Immunohistochemistry

Cryosections were washed with PBS and blocked for 1 h at room temperature with blocking solution (10% Lamb serum and 0.3% Triton X-100), and incubated overnight at 4°C with primary antibodies diluted in blocking solution. The following primary antibodies were used in this study: mouse anti-Ki67 (1:500; BD Biosciences, 550609); rabbit anti-Sox2 (1:1000; Abcam, ab92494); rabbit anti-Tbr2 (1:1000; Abcam, ab23345); rabbit anti-DCX (1:1000; Abcam, ab18723); rat anti-GFAP (1:500; Zymed, 13-300); rat anti-RFP (1:1000, Chromotek, 5f8-100); rat anti-BrdU (for BrdU or CldU detection) (1:500, Abcam, ab6326) and mouse anti-BrdU (for IdU detection)(1:100, BD Biosciences, 347580). For staining of Ki67, Tbr2, Sox2 and thymidine analogs, sections were heated in 10 mM Citric acid pH 6.0 on boiling water bath for 10 min prior to blocking. After three washes in PBS, sections were incubated for 2 h with corresponding secondary antibodies. After a final rinse with PBS, sections were mounted on glass slides with Prolong gold antifade reagent (Thermo Fisher Scientific).

#### Cell counting

Images were acquired using a Zeiss LSM 780 confocal microscope or Axio Scan Z.1 (Carl Zeiss). NIH ImageJ was used to count the cells. Brain sections were serially mounted on individual fifteen slides glasses. Cell counting was performed on every fifteenth sections containing DG at the same anatomical level between each group, and marker-positive cells were counted in the series of collected sections throughout the indicated areas in the DG. The total number of marker-positive cells in each indicated areas was obtained by multiplying the resultant counts by 15 (according to the interval between sections).

#### Real-time PCR

Total RNA was isolated using TRIzol™ Reagent (Thermo Fishire Scientific), according to the manufacturer’s instructions, and each sample was reverse-transcribed using a SuperScript IV cDNA Synthesis Kit (Invitrogen). Quantitative PCR reactions were performed using a KAPA SYBR Fast qPCR Kit (KAPA Biosystems) with ROX as reference dye, and transcript expression was measured via Applied Biosystem 7500 Real-Time PCR System (Life Technologies). Expression levels of each gene were normalized to RNA polymerase II subunit A (polr2a) and calculated relative to the control. Following primers were used for this study: Gli1 Fw: CCGACGGAGGTCTCTTTGTC; Gli1 Rv AACATGGCGTCTCAGGGAAG; Gli2 Fw: TGAAGGATTCCTGCTCGTG; Gli2 Rv: GAAGTTTTCCAGGACAGAACCA; Gli3 Fw: AAGCGGTCCAAGATCAAGC; Gli3 Rv: TTGTTCCTTCCGGCTGTTC; Polr2a Fw: CATCAAGAGAGTGCAGTTCG; Polr2a Rv: CCATTAGTCCCCCAAGTTTG.

#### Statistical analysis

At least three mice per group were analyzed. Statistical analyses were performed using either Student’s t-test (for comparisons between two groups) or one-way ANOVA with Tukey’s multiple comparison test (for multiple groups comparison) with Prism software (Graphpad). All experiments were independently replicated at least three times. Differences were considered statistically significant at p < 0.05. Asterisks indicate significant differences (* < 0.05; ** < 0.01, *** < 0.001, **** < 0.0001).

## Results

### Deletion of *Sufu* in NSCs reduces Shh signaling during DG development

Sufu is expressed in NSCs in the developing forebrain including presumptive DG cells (Yabut et al., 2015). We previously reported that deletion of *Sufu* in NSCs at early gestational stages (E10.5) severely disrupted the overall cytoarchitecture of the forebrain as a consequence of ectopic activation of Shh signaling (Yabut et al., 2015). To determine the effects of Sufu specifically in DG development, we used a hGFAP-Cre line to delete *Sufu* in NSCs at E13.5, before the initiation of DG development (E14.5) – we call these mice hGFAP-Sufu-KO. We first asked if deletion of *Sufu* increases Shh signaling activity by assessing the distribution of Shh-responding cells in the developing DG of hGFAP-Sufu-KO;Gli1^LacZ/+^ mice, which carry the Hh-responding transgene, Gli1-LacZ. Shh ligands originate from amygdala neurons and the adjacent ventral dentate neuroepithelium to activate Shh signaling in ventral hippocampal NSCs (Li et al., 2013). These Shh-responding NSCs subsequently migrate to the dorsal DG and gradually accumulate between the hilus and GCL to form the SGZ postnatally (Li et al., 2013). Accordingly, we found abundant Gli1-lacZ+ cells in the ventricular zone of the ventral hippocampus in Sufu^fl/fl^;Gli1^lacZ/+^ mice (Figure 1A). Gli1-lacZ+ cells were also present in the dorsal DG of Sufu^fl/fl^;Gli1^lacZ/+^ mice at P0 and were enriched in the SGZ at P7 (Figure 1B-C). Surprisingly, we found a remarkable reduction of Gli1-LacZ+ cells in ventral hippocampus of hGFAP-Sufu-KO;Gli1^lacZ/+^ mice at E15.4-P0, and small numbers detected throughout the anterior to posterior DG at P7. These data demonstrate that deletion of *Sufu* in NSCs decreases Shh signaling activity during DG development. This is in distinction to the situation in the neocortex, where deletion of Sufu increases Shh signaling acitivity (Yabut et al., 2015).

**Figure 1.**
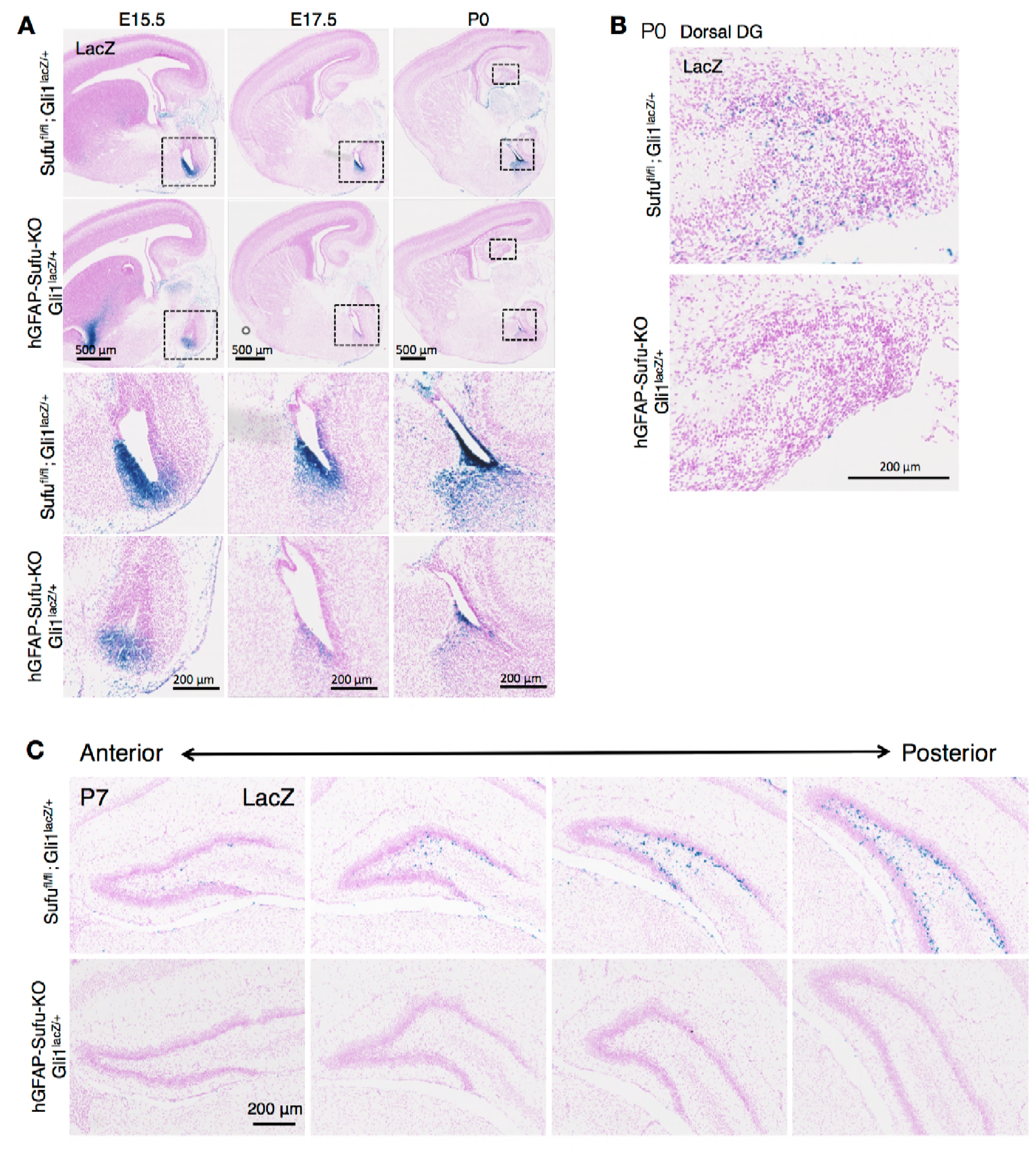
Deletion of *Sufu* decreases Hh-respoding cells during DG development. (A) Representative Gli1-LacZ staining images of sagittal brain sections in Sufu^fl/fl^;Gli1^lacZ/+^ and hGFAP-Sufu-KO;Gli1^lacZ/+^ mice at E15.5-P0. Magnified images of the black dashed-line boxes are shown in the lower two rows. (B) Representative Gli1-LacZ staining images of dorsal DG in sagittal sections of Sufu^fl/fl^;Gli1^lacZ/+^ and hGFAP-Sufu-KO;Gli1^lacZ/+^ mice. (C) From anterior to posterior, four levels of coronal sections for Gli1-nLacZ staining at E17.5 are shown. Note that lacZ+ cells are diminished in the DG of hGFAP-Sufu-KO;Gli1^lacZ/+^ mice from the beginning of DG development.

### Deletion of Sufu decreases proliferation of NSCs in developing DG

Shh signaling plays a pivotal role in establishing and maintaining the NSC pool to adulthood (Choe et al., 2015; Han et al., 2008; Li et al., 2013). Ablation of Shh signaling in NSCs by deleting *Smo* leads to a drastic reduction in NSC proliferation and results in a failure of SGZ establishment (Han et al., 2008; Li et al., 2013). Since hGFAP-Sufu-KO;Gli1^lacZ/+^ mice showed a reduction of Gli1 expression at the onset of DG development, we investigated if deletion of *Sufu* influences the proliferation capacity of NSCs particularly in the SGZ where Sox2+ cells form the NSC pool (Figure 2A). Although we found no difference in the number of Sox2+ cells between Sufu^fl/fl^ mice and hGFAP-Sufu-KO mice in the SGZ (Figure 2B), there was a significant reduction in Ki67+ proliferating cells in the P7 hGFAP-Sufu-KO mice (Figure 2C). In addition, there was a significant reduction in ratio of Ki67+ cells to Sox2+ NSCs in the SGZ of hGFAP-Sufu-KO mice (Figure 3D) indicating that very few Sox2+ cells retained their proliferative capacity. This contrasts starkly with the phenotypes seen in the *hGFAP-Cre;SmoM2* mice, in which Shh signaling is constitutively active in NSCs, where the number of Ki67+ cells in the SGZ and the proliferating population of Sox2+ cells were significantly higher than WT mice (Figure 2E-H). Taken together, these findings establish the crucial roles of Shh signaling in regulating NSC proliferation in the developing DG. Importantly, that deletion of *Sufu* resulted in downregulation, instead of ectopic activation, of Shh signaling in DG NSCs, indicates the distinct effects of Sufu in specific NSC subtypes and may involve a novel mechanism by which Sufu controls Shh signaling activity in DG NSCs.

**Figure 2.**
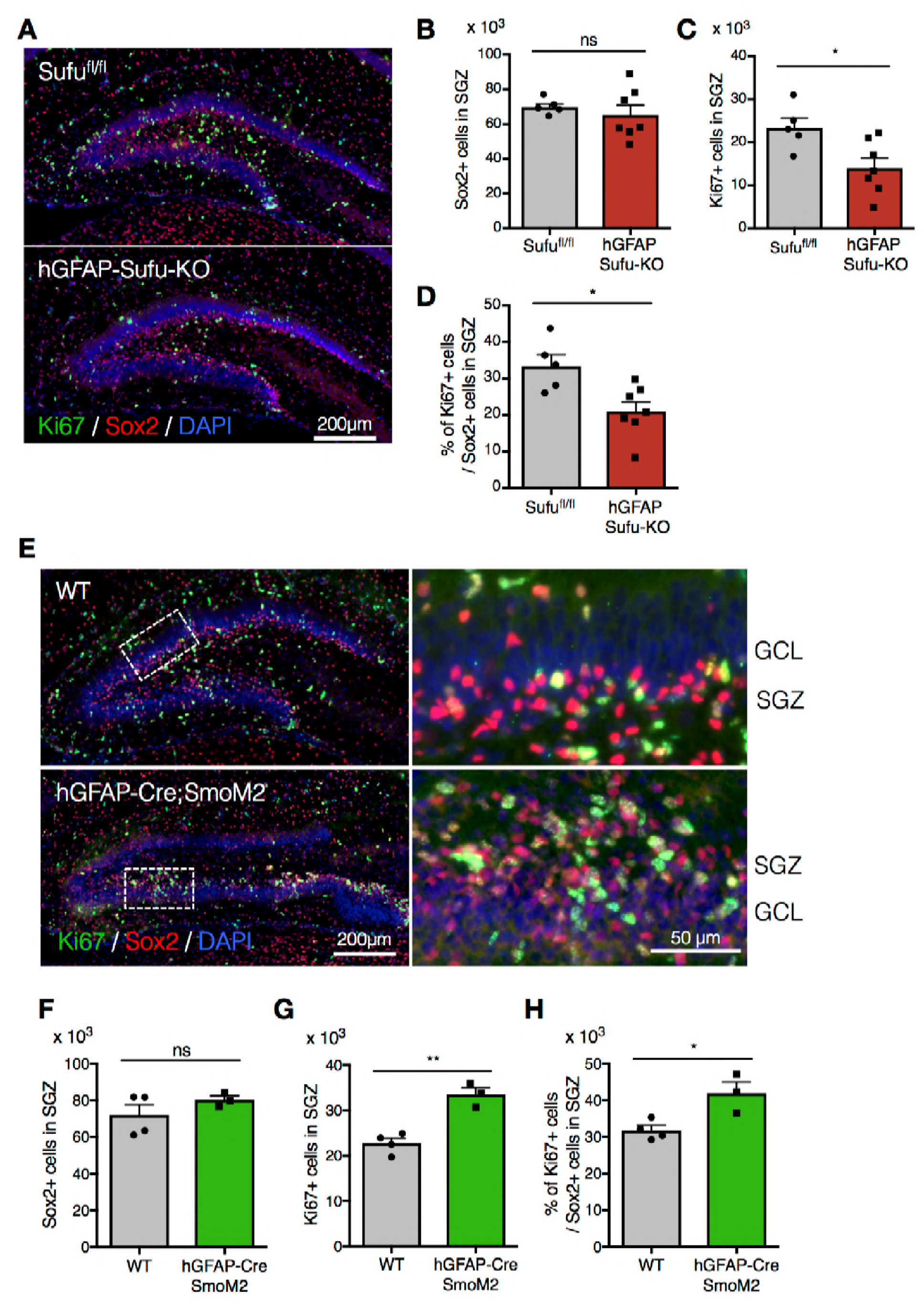
Deletion of *Sufu* decreases proliferating cell population in DG-NSCs. (A) Representative immunofluorescence images of Ki67 (green) and Sox2 (red) in the DG of *Sufu^fl/fl^* and hGFAP-Sufu-KO mice at P7. DNA is stained with DAPI (blue). (B,C) Quantification of Sox2+ (B) and Ki67+ (C) cells in the SGZ of *Sufu^fl/fl^* and hGFAP-Sufu-KO mice. Note that Ki67+ proliferating cells and proliferating NSCs population in SGZ are decreased in hGFAP-Sufu-KO mice. (D) The bar graph indicates the ratio of Ki67+ cells to Sox2+ NSCs in DGs of *Sufu^fl/fl^* and hGFAP-Sufu-KO mice. (E) Representative immunofluorescence images of Ki67 (green), Sox2 (red) and DAPI (blue) in the DG of WT and *hGFAP-Cre;SmoM2* mice at P7. Magnified images of the white dashed-line boxes are shown to the right of each image. (E,F) Quantification of Sox2+ (E) and Ki67+ (F) cells in the SGZ of WT and *hGFAP-Cre;SmoM2* mice. (G) The bar graph indicates the ratio of Ki67+ cells to Sox2+ NSCs in DGs of WT and *hGFAP-Cre;SmoM2* mice. Note that *hGFAP-Cre;SmoM2* mice shows opposite phenotype to hGFAP-Sufu-KO mice. Values represent mean ± SEM; NS: P > 0.05, *P < 0.05, **P < 0.01. Student’s *t*-test.

### In the absence of Sufu, Gli1 function becomes responsible for proper proliferation of NSCs during DG development

Gli1, Gli2 and Gli3 are the main transcription factors that transduce Shh signaling to downstream targets. Gli3 mainly functions as a gene repressor (Hu et al., 2006; Litingtung et al., 2002; Persson et al., 2002; Wang et al., 2014), whereas Gli1 and Gli2 are responsible for activating target gene expression (Bai and Joyner, 2001; Park et al., 2000). Gli2 and Gli3 play major roles in regulating gene expression during forebrain development, whereas Gli1 is largely dispensable (Bai et al., 2002; Park et al., 2000). Indeed, in the developing DG, we did not observe any differences in the number of Sox2+ cells and proliferating cells in the SGZ between Sufu^fl/fl^, Sufu^fl/fl^;Gli1^lacZ/+^ and Sufu^fl/fl^;Gli1^lacZ/lacZ^ mice (Figure S1). Therefore, we decided to use those genotypes as control. However, deletion of one (hGFAP-Sufu-KO;Gli1^lacZ/+^) or both (hGFAP-Sufu-KO;Gli1^lacZ/lacZ^) Gli1 alleles in hGFAP-Sufu-KO mice led to a more profound phenotype. Ki67+ cells and the proliferating population of Sox2+ cells in the SGZ were significantly reduced in hGFAP-Sufu-KO;Gli1^lacZ/+^ and hGFAP-Sufu-KO;Gli1^lacZ/lacZ^ mice compared with control (Figure 3A-C). In addition, we observed changes in NSC populations; hGFAP-Sufu-KO;Gli1^lacZ/+^ and hGFAP-Sufu-KO;Gli1^lacZ/lacZ^ mice displayed a remarkable decline in Sox2+ cells in the SGZ in contrast with KO mice, (Figure 3D). The number of Tbr2+ neuronal precursor cells, which was significantly reduced in KO mice, was further reduced in hGFAP-Sufu-KO;Gli1^lacZ/+^ mice (Figure 3E,F). This suggests that deletion of *Sufu* decreases the neurogenic competence of NSCs, and this worsens when *Gli1* expression is reduced.

**Figure 3.**
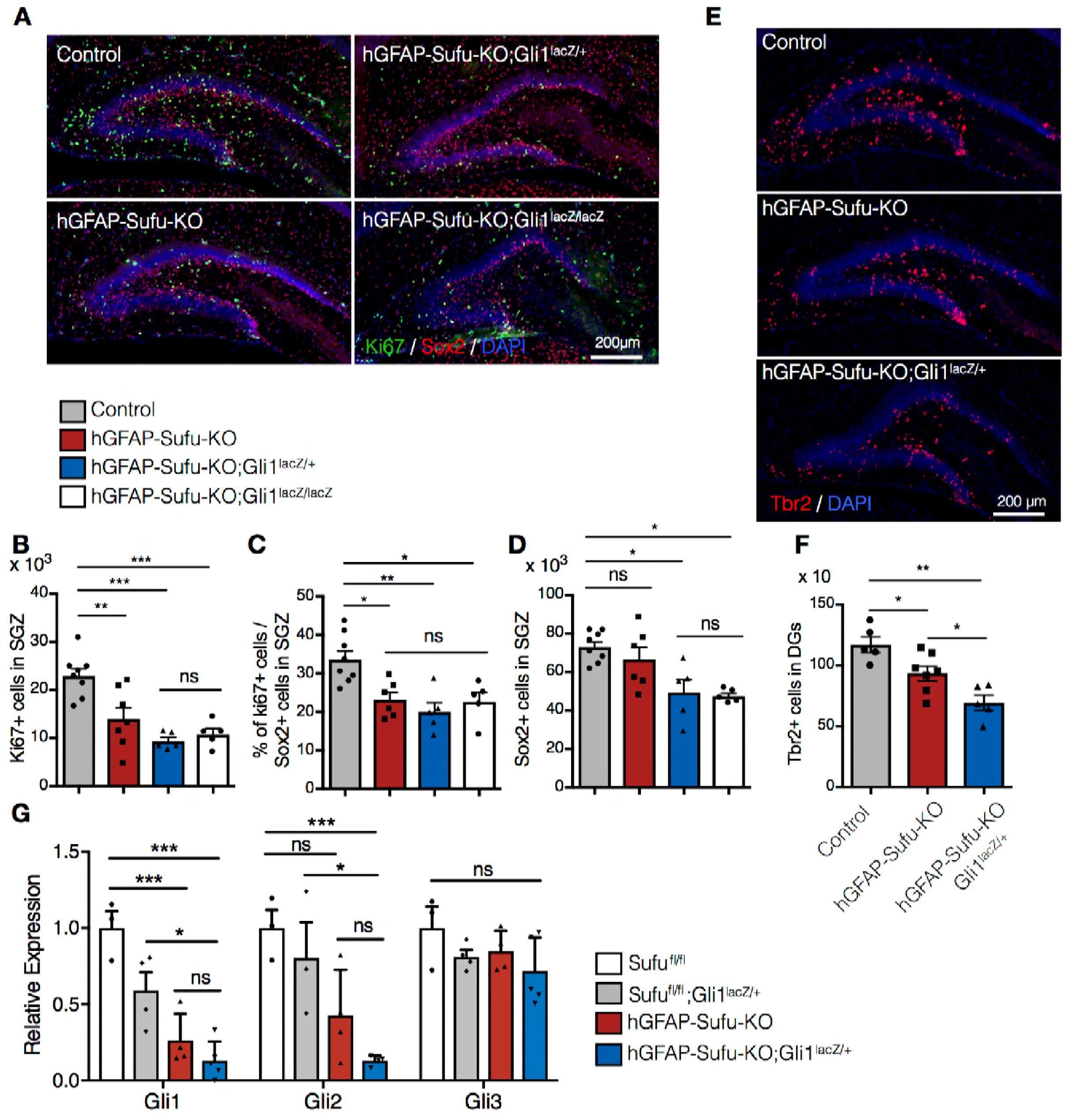
*Gli1* deletion increases the developmental defects in hGFAP-Cre Sufu^fl/fl^ mice. (A) Representative immunofluorescence images of Ki67 (green), Sox2 (red) and DAPI (blue) in the DG of Control, hGFAP-Sufu-KO, hGFAP-Sufu-KO;Gli1^lacZ/+^, and hGFAP-Sufu-KO;Gli1^lacZ/lacZ^ mice at P7. (B, C) Quantification of Ki67+ cells (B) and ratio of Ki67+ Proliferating cells population in Sox2+ cells (C) in SGZ. (D) Sox2+ cells counts in SGZ from each group. (E) Representative immunofluorescence images of Tbr2 (red) and DAPI (blue) in the DG of Control, hGFAP-Sufu-KO and hGFAP-Sufu-KO;Gli1^lacZ/+^ mice at P7. (E) The number of Tbr2+ cells in DGs. Adding *Gli1* deletion in absence of Sufu aggravates the *Sufu* deletion phenotypes. (F) qRT-PCR analyses of Gli1, Gli2 and Gli3 expression in the DGs of *Sufu^fl/fl^*, *Sufu^fl/fl^;Gli1^lacZ/+^*, hGFAP-Sufu-KO and hGFAP-Sufu-KO;Gli1^lacZ/+^ mice at P0. Expression of Shh signaling activator, Gli1 and Gli2, are decreased in hGFAP-Sufu-KO;Gli1^lacZ/+^ mice. Values represent mean ± SEM; NS: P > 0.05, *P < 0.05, **P < 0.01, ***P < 0.001. ANOVA with Tukey post-hoc tests.

Earlier, we showed a reduction in Shh signaling activity in hGFAP-Sufu-KO mice by lacZ staining (Figure 1). To further investigate Shh signaling activity in hGFAP-Sufu-KO mice, we extracted RNA from P0 DGs and examined the expression of *Gli1*, *Gli2* and *Gli3* by qPCR. Similar to LacZ staining results, *Gli1* expression was significantly down-regulated in hGFAP-Sufu-KO mice, indicating that deletion of *Sufu* decreases Shh signaling activity (Figure 3G). Interestingly, hGFAP-Sufu-KO;Gli1^lacZ/+^ mice showed remarkable reduction in *Gli2* expression in addition to *Gli1* expression. Both Gli activators were down-regulated in the absence of Sufu, suggesting that reducing *Gli1* expression in *Sufu* deletion further decreases Shh signaling activity in NSCs. Taken together, these data suggest that *Sufu* deletion increases the dependency on Gli1 function during DG development, and that proliferation of NSCs is severely impaired when *Gli1* expression is reduced in absence of Sufu.

### Deletion of *Sufu* during DG development decreases NSC number and impairs adult neurogenesis

After development, NSCs in the DG are maintained until adulthood and produce neurons throughout life in the rodent brain. Given the widespread impairments in NSC numbers and proliferative capacity in the absence of Sufu, we next investigated the impact of these defects on adult neurogenesis. To label newborn neurons in adult mice, we administrated the thymidine analog 5-bromo-2’-deoxyuridine (BrdU) to 8-week old mice for 5 days and sacrificed the animal 3 days post-BrdU injection (Figure 4A). In hGFAP-Sufu-KO;Gli1^lacZ/+^ mice, the number of newborn neurons decreased, reflecting the degree of proliferation impairments between hGFAP-Sufu-KO and hGFAP-Sufu-KO;Gli1^lacZ/+^ mice during development. To clarify the reason for reduced neurogenesis in hGFAP-Sufu-KO;Gli1^lacZ/+^ mice, we compared neurogenic competence in adult NSCs by calculating the number of newborn neurons produced from BrdU labeled cells, and found that there was no difference in the ratio of DCX with BrdU between three groups (Figure 4D), suggesting that deletion of *Sufu* at developmental stages did not affect the neurogenic competence of NSCs in adult DGs. This prompted us to investigate whether reduction of newborn neurons in hGFAP-Sufu-KO;Gli1^lacZ/+^ mice results from reduced NSC pool. Adult NSCs are maintained in a quiescent state until stimulated to proliferate and produce neurons (Encinas et al., 2011; Mira et al., 2010). These quiescent NSCs display a radial morphology with fiber extending to the molecular layer (Bignami and Dahl, 1974; Lugert et al., 2010; Rickmann et al., 1987; Sievers et al., 1992). To determine the number of quiescent NSCs in adult DGs, we counted the number of Sox2+ cells in SGZ, which have a GFAP+ radial fiber. The number of Sox2+/GFAP+ radial NSCs was significantly reduced in both hGFAP-Sufu-KO and hGFAP-Sufu-KO;Gli1^lacZ/+^ mice, with a greater reduction observed in hGFAP-Sufu-KO;Gli1^lacZ/+^ mice (Figure 4E,F). These data suggest that deletion of *Sufu* during developing DGs decreases the number of NSCs maintained through adulthood resulting in impaired adult neurogenesis.

**Figure 4.**
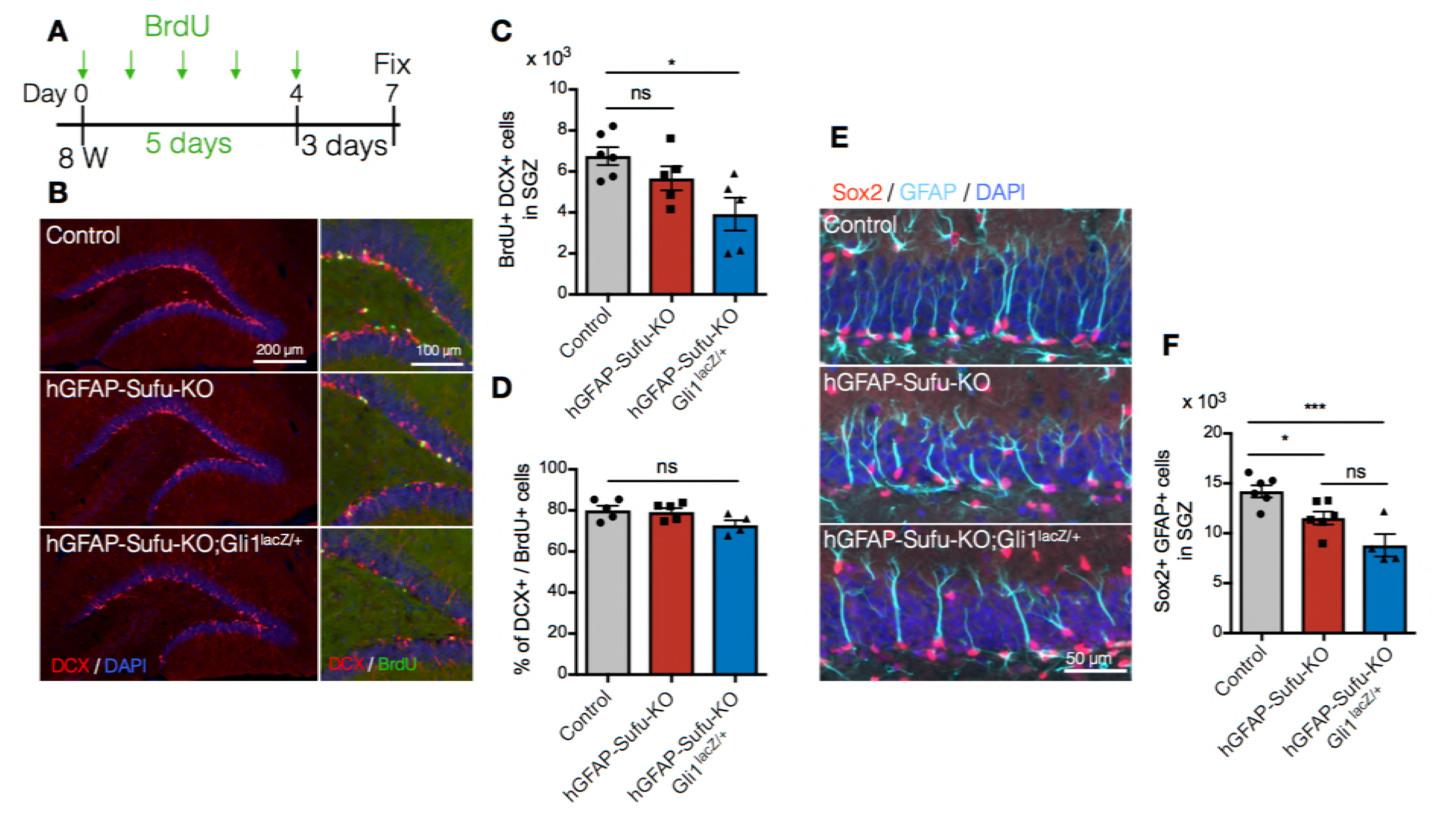
Deletion of *Sufu* during DG development decreases qNSCs pool in adult DGs. (A) Experimental scheme of BrdU injection. 8 week-old mice were injected with BrdU for 5 days and analyzed 3 days after last BrdU injection. (B) Representative immunofluorescence images for DCX (red) and DAPI (blue) in the DGs of control, hGFAP-Sufu-KO and hGFAP-Sufu-KO;Gli1^lacZ/+^ mice. Magnified images of DCX (red), BrdU (green) and DAPI (blue) are shown to the right of each image. (C) Quantification of BrdU+/DCX+ cells in SGZ. (D) The bar graph indicates the ratio of DCX+ cells to BrdU+ cells in SGZ. Representative immunofluorescence images for Sox2 (red), GFAP (cyan) and DAPI (blue) in the SGZ of control, hGFAP-Sufu-KO and hGFAP-Sufu-KO;Gli1^lacZ/+^ mice. (F) Quantification of qNSCs, indicated by radial GFAP+ fiber and Sox2 expression. Values represent mean ± SEM; NS: P > 0.05, *P < 0.05, ***P < 0.001. ANOVA with Tukey post-hoc tests.

### Deletion of *Sufu* impairs proliferation and expansion of NSCs in the DG at early postnatal stages

Proliferation of NSCs in the first postnatal week is critical for producing and maintaining NSCs until adult stages (Youssef et al., 2018). Elimination of proliferating cells in the first postnatal week, but not at 2-3 weeks, severely impairs the size of the NSC pool in adult DG. We have previously shown that long-lived NSCs of the DG are composed of Hh-responding cells in the ventral hippocampus at E17.5 (Li et al., 2013). Using Gli1^CreER/+^::Rosa^Ai14/+^mice treated with Tamoxifen at E17.5, we labeled Hh-responding cells and traced their migration during the first postnatal week. We found that Ai14+ cells were sparsely localized in the dorsal DG at P0 (Figure 5A). However, the number of Ai14+ cells dramatically increased with development and accumulated in the border between hilus and GCL from P3 to P7. The number of Sox2+ cells in Ai14+ cells of ventrally derived NSCs was significantly increased from P0 to P3, and P3-P7, respectively (Figure 5B-D). This suggests that the first postnatal week is a critical period for long-lived NSC expansion.

**Figure 5.**
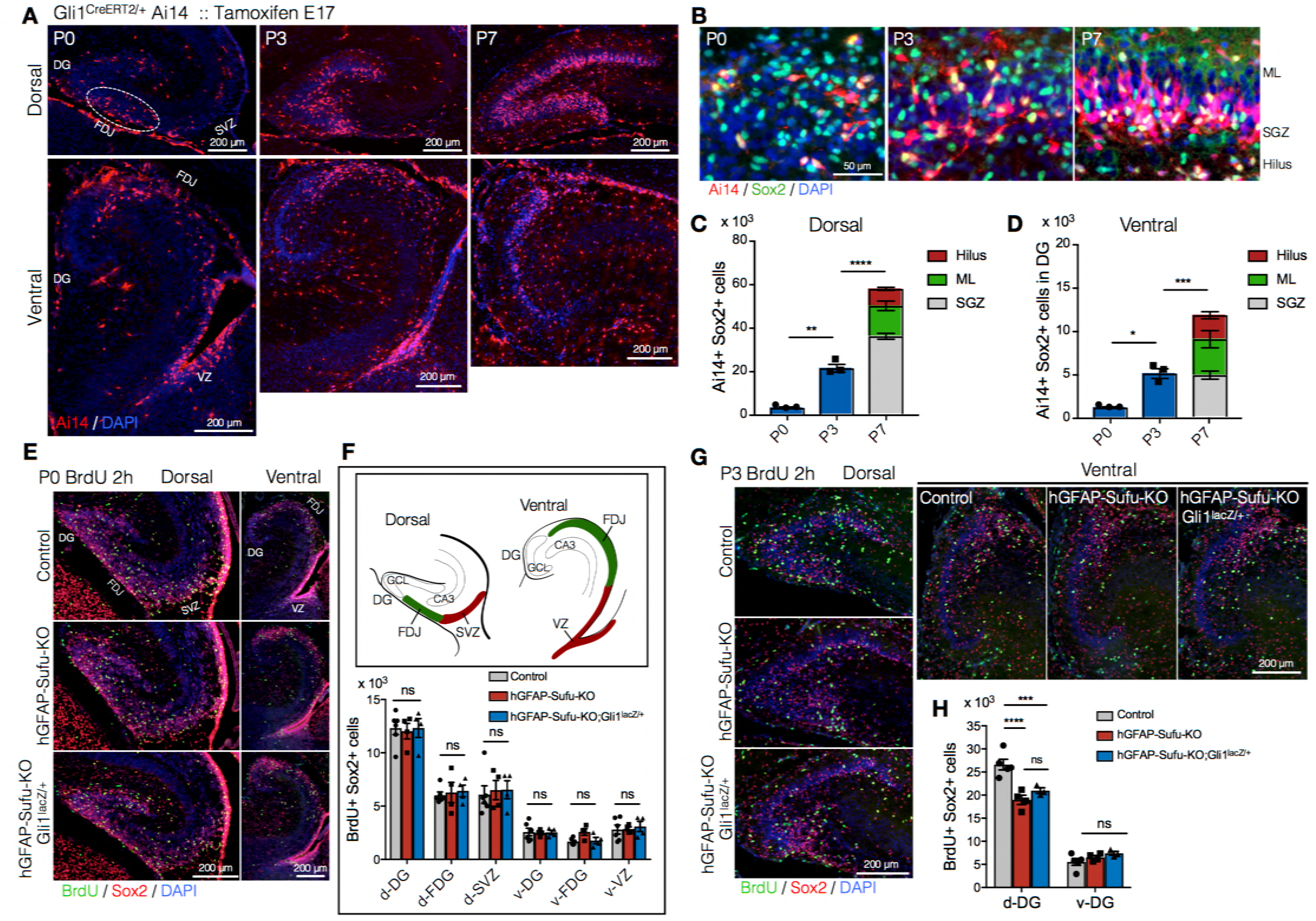
Long-lived NSCs expand in first postnatal week, and loss of Sufu impairs its expansion. (A) Fate tracing of Hh-responding cells at E17.5 in postnatal DG development. Gli1^CreERT2/+^ Ai14/+ mice were treated with tamoxifen at E17.5 and Ai14+ Hh-responding cells are analyzed in postnatal first week (P0-7). Representative immunofluorescence images for Ai14 (red) and DAPI (blue) in dorsal and ventral hippocampus in sagittal sections. Note that Hh-responding cells labeled by Ai14 at E17.5 sparsely appear in DGs at P0 and subsequently accumulate in SGZ. (B) Representative immunofluorescence images for Ai14 (red), Sox2 (green) and DAPI (blue) in SGZ. (C,D) Quantification of Ai14+/Sox2 + cells in dorsal (C) and ventral (D) DGs. The number of long-lived NSCs, indicated as Ai14+/Sox2+ cells, are increased in postnatal first week. (E) BrdU was injected at P0, and pups were sacrificed 2h later. Representative immunofluorescence images for BrdU (green), Sox2 (red) and DAPI (blue) in dorsal and ventral DGs of control, hGFAP-Sufu-KO and hGFAP-Sufu-KO;Gli1^lacZ/+^ mice. (F) Schematic illustration of the dorsal and ventral DG at P0. d: dorsal, v: ventral, FDJ: fimbriodentate junction; SVZ: subventricular zone. The bar graph indicates the number of Sox2+/BrdU+ cells in each region. (G) BrdU was injected at P3, and pups were sacrificed 2h later. Representative immunofluorescence images for BrdU (green), Sox2 (red) and DAPI (blue) in dorsal and ventral DGs of control, hGFAP-Sufu-KO and hGFAP-Sufu-KO;Gli1^lacZ/+^ mice. (H) Quantification of Sox2+/BrdU+ cells in dorsal and ventral DGs. Values represent mean ± SEM; NS: P > 0.05, *P < 0.05, **P < 0.01, ***P < 0.001, ****P < 0.0001. ANOVA with Tukey post-hoc tests.

In hGFAP-Sufu-KO mice, we found that the number of NSCs in adult DG was significantly reduced (Figure 4E,F). This might be attributed to the failure of NSC expansion in the first postnatal week. To address this possibility, we labeled proliferating cells with BrdU at P0 or P3, and assessed the number of BrdU+/Sox2+ cells 2 hours post BrdU injection. At P0, there was no difference in the number of BrdU+/Sox2+ cells in both dorsal and ventral DGs between control, hGFAP-Sufu-KO and hGFAP-Sufu-KO;Gli1^lacZ/+^ mice (Figure 5E,F). However, at P3, in dorsal, but not ventral DG, both hGFAP-Sufu-KO and hGFAP-Sufu-KO;Gli1^lacZ/+^ mice showed significant reduction in BrdU+/Sox2+ cells compared with control (Figure 5G,H). These findings showed that Sox2+ NSCs in the dorsal DG remain proliferative during the expansion period of long-lived NSCs. Taken together, these data suggest that *Sufu* deletion impairs the proliferation of NSCs at a critical expansion period resulting in reduced number of quiescent NSCs in the adult DG.

### Deletion of *Sufu* leads to the premature transition of NSCs into quiescence during DG development

Our data show that *Sufu* deletion decreased NSC proliferation during the critical expansion period for long-lived NSCs, pointing to the likelihood that NSCs prematurely transitioned into a quiescent state. To test this, we utilized two thymidine analogs 5-Chloro-2-deoxyuridine (CldU) and 5-Iodo-2-deoxyuridine (IdU), and injected each thymidine analogs at different time points; CldU at P0, 3, 7 or 14 and IdU at 8 weeks old (Figure 6A,B). Because the thymidine analog is diluted as cells divide, cells that proliferated and stopped in the postnatal period will have detectable CldU, and therefore, when IdU is injected at adult stages, cells that became quiescent in developmental stages and then are reactivated in the adult will be double positive for CldU and IdU in adult stages. Double positive cells were observed more in mice injected with CldU at P3 and gradually decreased in groups injected with CldU at later postnatal stages (Figure 6C,D). Accordingly, the number of DCX+/CldU+/IdU+ cells in newborn neurons was highest in groups injected at P3 (Figure 6E,F). Similarly, control mice injected with CldU at P3 had the highest number of CldU and IdU double positive cells at adult stages (Figure 6G,H). These data suggest that around P3-P7, NSCs in the control DG reduce their proliferation rate and become quiescent. However, in hGFAP-Sufu-KO and hGFAP-Sufu-KO;Gli1^lacZ/+^ mice, the number of CldU and IdU double positive cells in CldU-injected groups at P3 or P7 was significantly decreased compared to control mice. Instead, detection of CldU and IdU double positive cells were significantly increased at P0 compared to control. The number of DCX+ cells double labeled with CldU and IdU was also decreased in P3 or P7 CldU injected groups, whereas it was increased significantly in P0 CldU injected groups (Figure 6I). These data suggest that NSCs in hGFAP-Sufu-KO and hGFAP-Sufu-KO;Gli1^lacZ/+^ mice prematurely exited the proliferative state from P0-3 instead of at P3-P7. In line with these observations, we found that the Ki67+ proliferating population in Sox2+ NSCs was significantly reduced at P3, but not P0 in the DGs of hGFAP-Sufu-KO and hGFAP-Sufu-KO;Gli1^lacZ/+^ mice compared with control (Figure S2). Taken together, these data suggest that deletion of *Sufu* precociously reduces proliferation of NSCs and leads to premature transition to quiescent state and thus a smaller pool of quiescent NSCs in the adult dentate.

**Figure 6.**
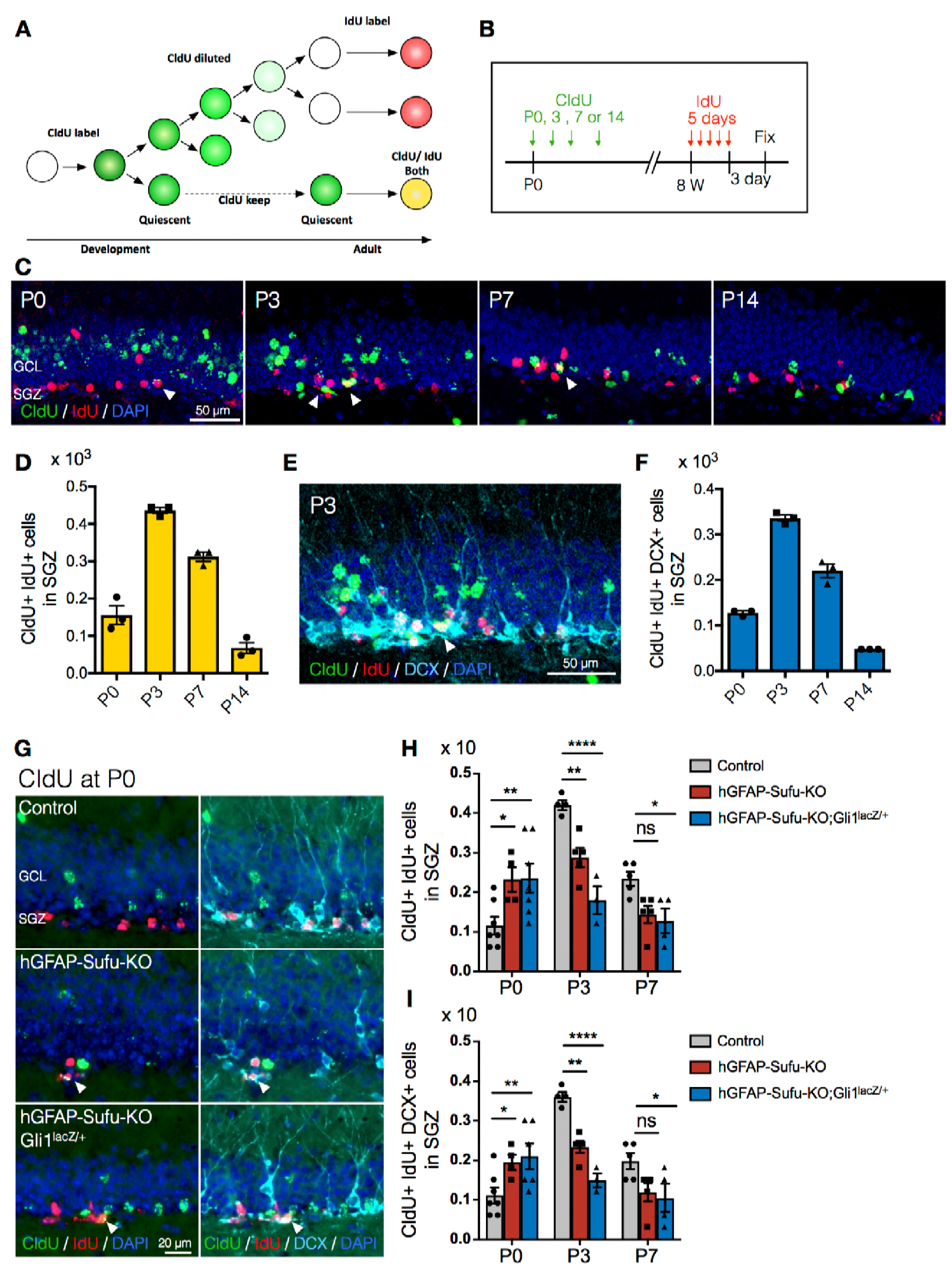
Deletion of *Sufu* prematurely induces the quiescent state transition. (A) Schematic illustration for labeling NSCs that established quiescent state during DG development using two thymidine analogs. CldU is diluted when cell divides. Thus CldU amount goes down if the cell continuously proliferates during DG development. However, if the cell becomes quiescent state, CldU amount is maintained until adult then the cell can be double-labeled with CldU and IdU if the CldU-labeled quiescent cell starts proliferation and incorporate IdU at adult. (B) Experimental scheme of CldU and IdU injection. CldU was injected at P0, P3, P7 or P14, and then IdU was injected for 5 days at 8 weeks old. The mice were sacrificed at 3 days after last IdU injection. (C) Representative immunofluorescence images for CldU (green), IdU (red) and DAPI (blue) in the SGZ of animals injected CldU at different stages. White arrowheads indicate the CldU/IdU double-labeled cells. (D) Quantification of CldU+/IdU+ cells in the SGZ. (E) Representative immunofluorescence images for CldU (green), IdU (red), DCX (cyan) and DAPI (blue) in the SGZ of animal injected CldU at P3. White arrowheads indicate the CldU+/IdU+/DCX+ cells. (F) Quantification of CldU+/IdU+/DCX+ cells in the SGZ. NSCs labeled with CldU at P3 maintained CldU until adult more than at other stages. (G) Representative immunofluorescence images for CldU (green), IdU (red), DCX (cyan) and DAPI (blue) in the SGZ of control, hGFAP-Sufu-KO and hGFAP-Sufu-KO;Gli1^lacZ/+^ mice injected CldU at P0. White arrowheads indicate the CldU+/IdU+/DCX+ cells. (H,I) Quantification of CldU+/IdU+ cells (H) and CldU+/IdU+/DCX+ (I) cells in the SGZ. In the absence of Sufu, NSCs labeled CldU at P0 more maintained CldU until adult. Values represent mean ± SEM; NS: P > 0.05, *P < 0.05, **P < 0.01, ****P < 0.0001. ANOVA with Tukey post-hoc tests.

## Discussion

Quiescence is key to maintaining the NSC pool and critical for enabling lifelong neurogenesis, so understanding how actively dividing NSCs become quiescent during development is vitally important. Here, we demonstrated that Sufu ensures the expansion of long-lived NSCs during postnatal DG development. We found that long-lived NSCs dramatically expand in the first postnatal week before entering the quiescent state over several days. *Sufu* deletion impairs the ability of long-lived NSCs to expand in the first postnatal week, which results in the premature entry of NSCs into the quiescent state (Figure 7). This defect is a result of decreased Shh signaling activity as a result of Sufu deletion in NSCs. Thus, Sufu modulates the timing of quiescence of NSCs in DG development by controlling NSCs expansion via modulation of Shh signaling activity.

**Figure 7.**
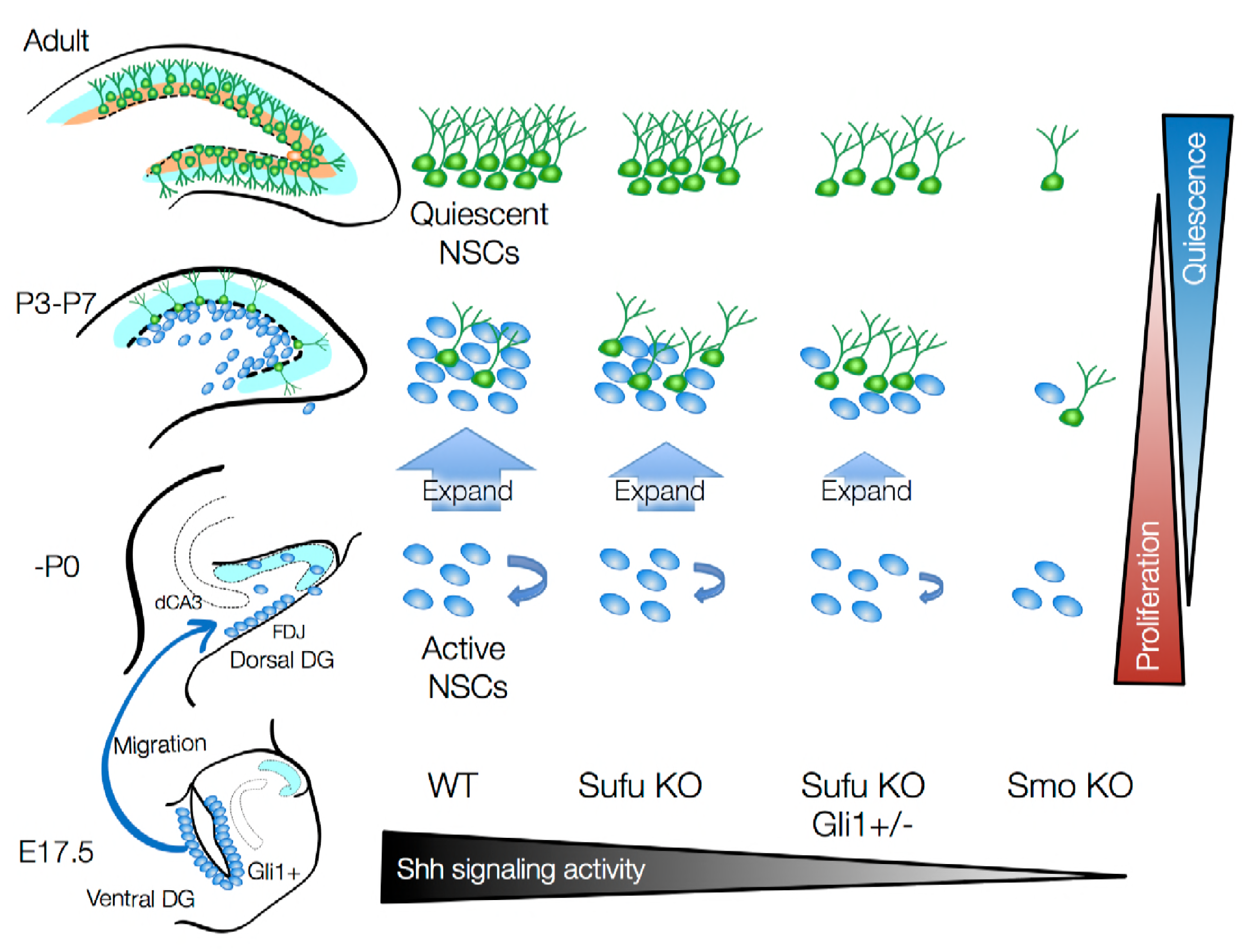
Sufu is important for the perinatal expansion and quiescent state transition of dentate NSCs. (A) Schematic summary illustrating the role of Sufu and Shh signaling activity in initial production, expansion and quiescent state transition of NSCs during DG development. Long-lived NSCs are produced from Hh-responding cells in ventral hippocampus by Shh stimuli at E17.5 and migrate to dorsal and ventral DGs. This subset of NSCs expands in first postnatal week and subsequently become quiescent state. Sufu controls shh signaling activity in NSCs during DG development. *Sufu* deletion decreases the Shh signaling activity and leads to impaired expansion of long-lived NSC, resulting in premature quiescent state transition and small NSC pool in adult. On the other hand, complete ablation of Shh signaling activity by deleting *Smo* impairs initial production of long-lived NSCs at the beginning of DG development and compromises the establishment of neurogenic niche.

Surprisingly, our data showed that deletion of *Sufu* in GFAP-expressing NSCs decreased Shh signaling activity. Gli1-LacZ+ cells were dramatically reduced in the embryonic ventral hippocampus and were abolished in the postnatal DG. Shh signaling is crucial for the initial production of dentate NSCs in the embryonic stage. *Smo* deletion in NSCs severely impairs initial NSC production and leads to decreased size of GCL (Han et al., 2008; Li et al., 2013). Although *Sufu* deletion impaired proliferation of NSCs in the first postnatal week, we still observed comparable number of proliferating NSCs at P0, indicating that initial NSC production at embryonic stages was not affected. Whether this process is dependent on Shh signaling is yet to be investigated, since we cannot exclude the possibility that very low levels of Shh signaling activity, likely undetectable by LacZ activity, are sufficient to produce the proper number of NSCs in embryonic stages.

The previously established roles of Sufu established it as a negative regulator of Shh signaling pathway through several mechanisms - promoting the formation of Gli repressors, removing Gli1 from nucleus, and recruiting transcription repressors to Gli-target genes sites (Barnfield et al., 2005; Cheng and Bishop, 2002; Kise et al., 2009; Kogerman et al., 1999). However, recent studies provide evidence that Sufu may have dual function enabling it to act as a negative and positive regulator of Shh signaling under specific conditions. For example, in mouse embryonic fibroblast (MEF), Shh ligand binding triggers Sufu translocation to the nucleus with Gli1, where it binds to and facilitates cytoplasmic export of Gli3 repressor, thereby enhancing Shh signaling activity (Zhang et al., 2017). Furthermore, increasing the amount of Sufu added into *Sufu^-/-^* MEF compromises Hh-responsiveness in the absence of exogenous Shh, which is consistent with negative function of Sufu for Shh signaling. However, in the presence of exogenous Shh, Hh-responsiveness of *Sufu^-/-^* MEF is dramatically elevated with the addition of increasing levels of Sufu, an effect that was not observed when Ptch1 was added (Chen et al., 2009). These reports demonstrate that Sufu can act to enhance or maximize Shh signaling activity. Considering these roles, decreased Shh signaling activity in Sufu KO mice could result from the failure to maximize Shh signaling activity through mechanisms that likely involve stabilization of Gli transcription factors. Supporting this, our data showed that *Gli1* and *Gli2* expression were down-regulated in hGFAP-Sufu-KO;Gli1^lacZ/+^ mice, whereas *Gli3* expression was not affected. Gli1 and Gli2 primarily act as activators of Shh-target gene expression while Gli3 mainly represses Shh target genes. In hGFAP-Sufu-KO;Gli1^lacZ/+^ mice, Gli3 repressor function may be dominant resulting in the reduced expression of Shh signaling activators and downregulation of Shh signaling activity in the absence of Sufu. Altogether, our data demonstrates multiple regulatory roles of Sufu in Shh signaling pathway that is dependent on cell type and context.

We found that deletion of *Sufu* increased the dependency of proper DG development on Gli1 function. Gli1 is not necessary for initial activation of Shh signaling (Bai et al., 2002). Therefore, *Gli1* deletion normally does not cause any significant developmental defects (Park et al., 2000). However, the phenotypes in hGFAP-Sufu-KO mice clearly worsen when combined with *Gli1* deletion, suggesting that Gli1 function becomes necessary in the absence of Sufu. Previous studies show that Sufu is important for the stabilization of Gli2 and Gli3 proteins (Makino et al., 2015; Wang et al., 2010). Loss of Sufu results in diminishing Gli2 full-length activator. Sufu competitively binds to Gli2 and Gli3 with speckle-type POZ protein (Spop), which recruits ubiquitin ligases and degrades the full-length forms of Gli2 and Gli3 (Chen et al., 2009; Wang et al., 2010; Wen et al., 2010). These finding provides a possibility that loss of Gli2 activators in the absence of Sufu increases the requirement for Gli1 function to maintain Shh signaling activity. Indeed, Gli1 is able to compensate for lost Gli2 activator function and rescue the developmental defects of Gli2 knockout mice (Bai and Joyner, 2001).

Expansion of long-lived NSCs must occur in the first postnatal week during which time ventral-derived NSCs dramatically increase their numbers in both the ventral and dorsal DG. Interestingly, *Sufu* deletion only impaired the proliferation of NSCs in the dorsal DG, but not ventral DG. This difference could be due to underlying molecular differences between NSCs and the surrounding cells residing in these regions. Linage tracing of Shh expressing cell shows that neurons in the medial entorhinal cortex and hilar mossy cells function as the local source of Shh ligand for ventral derived NSCs, while hair mossy cells in dorsal DG, but not ventral DG, are demonstrated as Shh sources (Li et al., 2013). Conditionally removing Shh ligand from these local neurons abolishes signaling in Hh-responding cells and results in reduction of NSCs number, indicating that local Shh ligands are important for the activation of Shh signaling in NSCs after migration to the dorsal DGs (Li et al., 2013). Similar to the mice with elimination of local Shh ligand, our data showed that deletion of *Sufu* abolished Hh-responding cells in postnatal DGs, followed by the reduction of proliferating NSCs in dorsal DGs. These findings indicate that dorsal and ventral NSCs use distinct developmental approaches to navigate the transition to produce long-lived NSCs. Dorsal NSCs must rely on Sufu to ensure optimal Shh signaling activity suitable for the expansion of long-lived NSCs during DG development, while for NSCs in the ventral DG appear able to expand and become quiescent without Sufu.

Our data suggests that Shh signaling activity must be continuously maintained to promote NSC expansion and that eventual reduction in Shh signaling activity promotes NSCs to transition into the quiescent state. Supporting this, we found that Gli1-LacZ+ cells were abundant in the ventral ventricular zone but progressively decreased after migration into the DG in the first postnatal week. Additionally, we found that constitutive activation of Shh signaling in NSCs, by conditional expression of SmoM2, increased proliferating NSCs and prevented transition into quiescence. These observations suggest that Sufu is important in sustaining NSC proliferation until NSCs begin to transition into a quiescent state. However, we also observed in SmoM2 mutants that some populations of NSCs become quiescent, indicating that reduction of Shh signaling activity alone is not sufficient to initiate quiescence and that other signaling mechanisms are involved in this process. Indeed, several extracellular factors can regulate the quiescent state of NSCs, such as Bone Morphogenetic Proteins (BMPs), Notch, and gamma-aminobutyric acid (GABA) (Kawaguchi et al., 2013; Mira et al., 2010; Song et al., 2012). These molecules are secreted by granule neurons, astrocytes, microglia, and interneurons in the DG and are likely sources of signals for migrating ventral-derived NSCs (Bonaguidi et al., 2011; Bond et al., 2014; Kawaguchi et al., 2013; Mira et al., 2010; Song et al., 2012; Yousef et al., 2014). The activity of these signaling pathways, and the simultaneous reduction in Shh signaling with development, may coordinately function to ensure successful transition of NSCs into the quiescent state during DG development.

## Acknowledgements

We thank members of the S.J.P. lab for helpful discussions, in particular O.R. Yabut and T.Bao for technical help, suggestions, and helping to write this manuscript. This research was supported by NIH grant R01 NS075188 (S.J.P.), Sasakawa Scientific Research Grant (H.N), The Uehara Memorial Foundation (H.N), JSPS Overseas Research Fellowships (H.N).

## Author contributions

H.N conceived of and performed experiments, analyzed the data, and wrote the manuscript. J.G.C, performed experiments and analyzed the data. K.N and S.J.P supervised, conceived of experiments, analyzed the data, and wrote the manuscript.

## Figure legends

**Supplemental figure 1.**
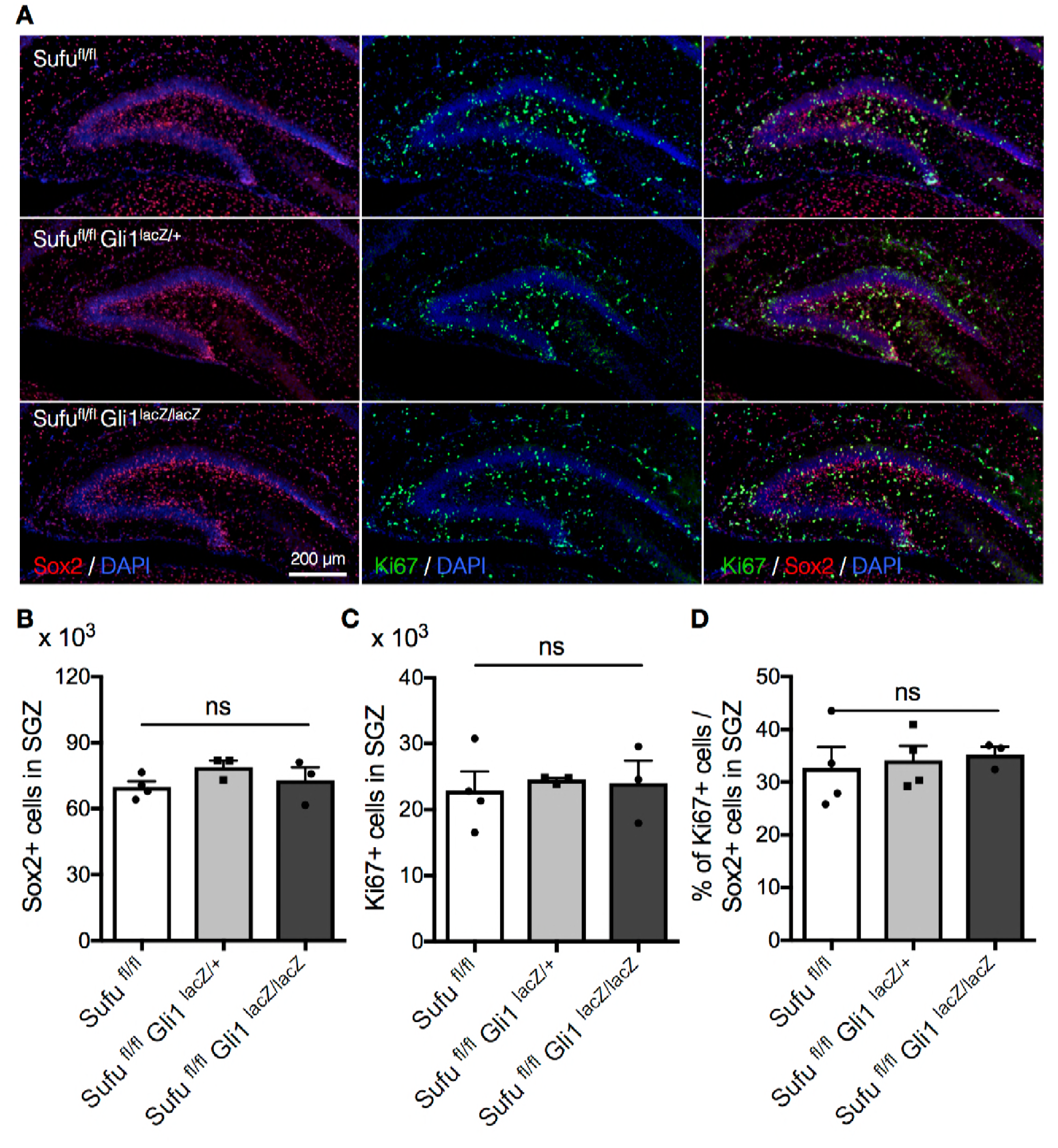
Deletion of *Gli1* does not affect cell proliferation. (A) Representative immunofluorescence images of Sox2 (red) and Ki67 (green) in the DG of *Sufu^fl/fl^*, *Sufu^fl/fl^*;*Gli1^lacZ/+^, Sufu^fl/fl^*;*Gli1^lacZ/lacZ^* mice at P7. DNA is stained with DAPI (blue). (B,C) Quantification of Sox2+ (B) and Ki67+ (C) cells in the SGZ. (D) The bar graph indicates the ratio of Ki67+ cells to Sox2+ NSCs in DGs. Values represent mean ± SEM; NS: P > 0.05. ANOVA with Tukey post-hoc tests.

**Supplemental figure 2.**
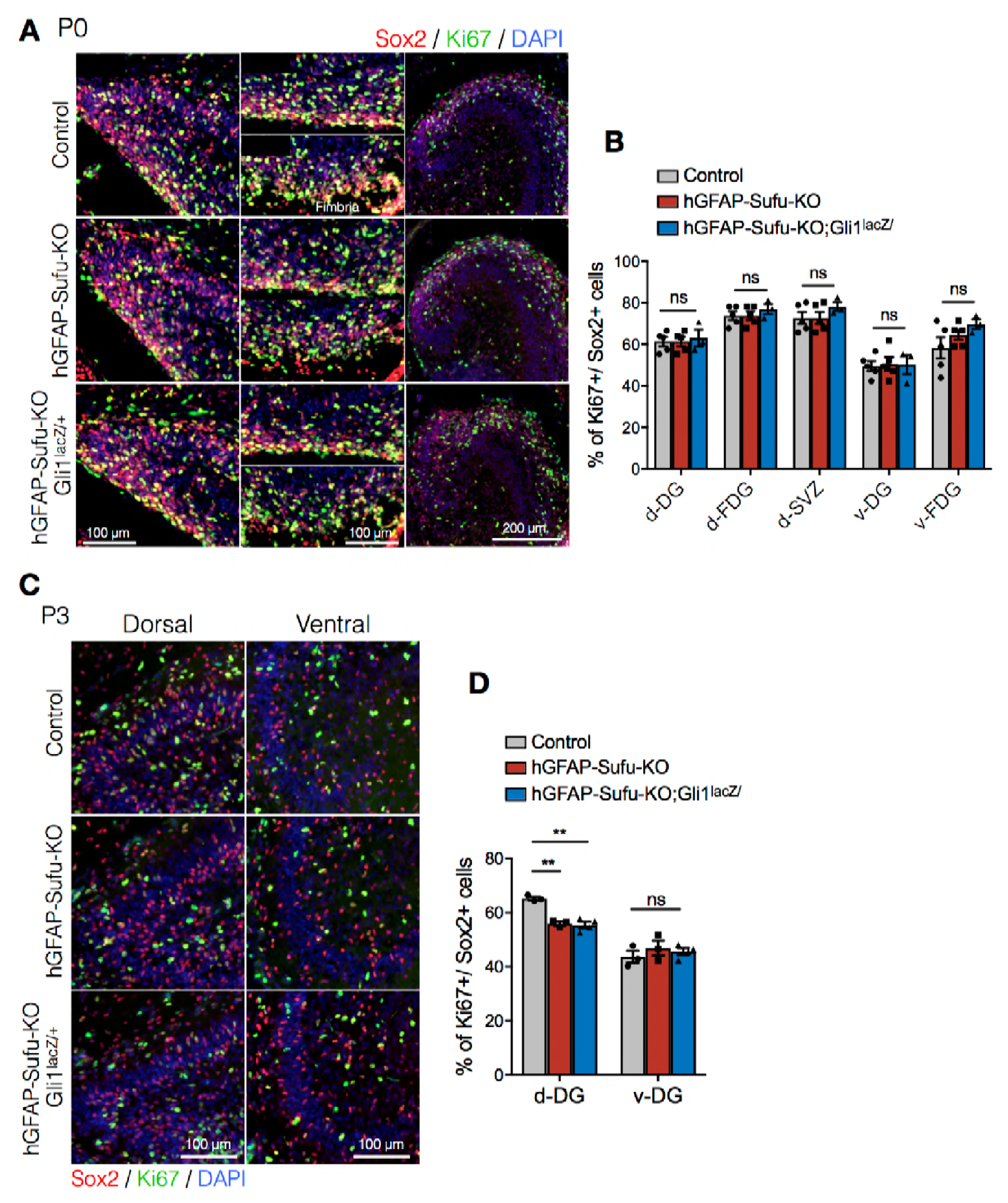
Deletion of *Sufu* decreases the proliferating cell population in Sox2+ NSCs at P3. (A) Representative immunofluorescence images of Sox2 (red) and Ki67 (green) in the dorsal and ventral DG of control, hGFAP-Sufu-KO and hGFAP-Sufu-KO;Gli1^lacZ/+^ mice at P0. DNA is stained with DAPI (blue). d: dorsal, v: ventral, FDJ: fimbriodentate junction; SVZ: subventricular zone. (B) The bar graph indicates the ratio of Ki67+ cells to Sox2+ NSCs in DGs. (C) Representative immunofluorescence images of Sox2 (red) and Ki67 (green) in the dorsal and ventral DG of control, hGFAP-Sufu-KO and hGFAP-Sufu-KO;Gli1^lacZ/+^ mice at P3. (D) The bar graph indicates the ratio of Ki67+ cells to Sox2+ NSCs in DGs. Values represent mean ± SEM; NS: P > 0.05, **P < 0.01. ANOVA with Tukey post-hoc tests.

